# Integration of optic flow into the sky compass network in the brain of the desert locust

**DOI:** 10.1101/2022.11.24.517785

**Authors:** Frederick Zittrell, Kathrin Pabst, Elena Carlomagno, Ronny Rosner, Uta Pegel, Dominik M. Endres, Uwe Homberg

## Abstract

Flexible orientation through any environment requires a sense of current relative heading that is updated based on self-motion. Global external cues originating from the sky or the earth’s magnetic field and local cues provide a reference frame for the sense of direction. Locally, optic flow may inform about turning maneuvers, travel speed and covered distance. The central complex in the insect brain is associated with orientation behavior and largely acts as a navigation center. Visual information from global celestial cues and local landmarks are integrated in the central complex to form an internal representation of current heading. However, it is largely unclear which neurons integrate optic flow in the central-complex network. We recorded intracellularly from neurons in the locust central complex while presenting lateral grating patterns that simulated translational and rotational motion to identify these sites of integration. Certain types of central-complex neurons were sensitive to visual self-motion independent of the type and direction of simulated motion. Columnar neurons innervating the noduli, paired central-complex substructures, were tuned to the direction of simulated horizontal turns. Modelling the connectivity of these neurons with a system of proposed compass neurons can account for rotation-direction specific shifts in the activity profile in the central complex corresponding to turn direction. Our model is similar but not identical to the mechanisms proposed for angular velocity integration in the navigation compass of the fly *Drosophila*.

## 1 INTRODUCTION

Animals navigate to feed, escape, migrate, and reproduce. Navigational tasks require a sense of current travel direction, which must be anchored to external cues and updated by internal cues, generated by ego-motion. Celestial cues are used as external cues by many insects, such as bees (von Frisch, 1946), ants (Fent, 1986), butterflies (Perez et al., 1997), dung beetles (Byrne et al., 2003), fruit flies (Weir and Dickinson, 2012), and caterpillars (Uemura et al., 2021). The sun and the skylight polarization pattern provide a reliable reference for dead reckoning (Gould, 1998). Internal cues, such as proprioceptive feedback (Wittlinger et al., 2006) and optic flow (Srinivasan, 2015; Stone et al., 2017), provide information about traveling speed and covered distance and may update the inner sense of direction in the absence of external cues. Only the flexible combination of information from external and internal cues enables robust and efficient navigation behavior, such as path integration (Heinze et al., 2018).

The central complex (CX), a midline spanning group of neuropils, houses the internal sense of direction in the brain of insects. It consists of the protocerebral bridge (PB), the lower (CBL) and upper (CBU) division of the central body, also termed ellipsoid- and fan-shaped body, and a pair of layered noduli (NO), and is associated with learning, memory and, importantly, spatial orientation (Pfeiffer and Homberg, 2014). The PB and the CBL are subdivided into series of 16 or 18 columns that are connected across the brain midline in a precise topographic manner (Pfeiffer and Homberg, 2014; Hulse et al., 2021; Hensgen et al., 2022).

CX neurons in various insect species are tuned to celestial cues (Heinze, 2017; Honkanen et al., 2019) and encode the solar azimuth in a compass-like manner in the locust (Pegel et al., 2019; Zittrell et al., 2020). Silencing compass neurons in the CX impairs navigation behavior in the fruit fly (Giraldo et al., 2018), showing the necessity of the CX for this behavior. Like mammalian head direction cells (Taube, 1998, 2007), specific CX neuron populations are tuned to the animal’s current heading (Seelig and Jayaraman, 2015; Hulse and Jayaraman, 2020). This internal heading estimate is multimodally tethered to environmental cues, such as visual compass cues and wind direction (Okubo et al., 2020), but also operates without external input, because internal cues from self motion are likewise integrated (Green et al., 2017; Turner-Evans et al., 2017; Green and Maimon, 2018).

Although the understanding of the CX network has made considerable progress, mainly owing to research in the fruit fly, and plausible models explaining network computations for navigation have been proposed (Stone et al., 2017), it is largely unclear at which network stages optic flow input is integrated in the sky compass network. To investigate this, we recorded intracellularly from various CX neurons in the desert locust (*Schistocerca gregaria*), a long-distance migratory insect, while stimulating laterally with wide-field visual motion that simulated self-motion to the animal. We analyzed general motion sensitivity for translational and rotational self-motion directions and tested whether the neural responses to opposing motion directions were discriminated (direction selectivity).

We implemented an algorithmic model (in the sense of Marr and Poggio (1979)) of the CX circuit which integrates visual self-motion cues with head direction representation. Modeling was guided by data on two types of columnar neurons with one being sensitive to the direction of simulated horizontal turns.

## 2 METHODS

### 2.1 Animals and preparation

Desert locusts (*Schistocerca gregaria*) were kept and dissected as described previously (Zittrell et al., 2020). Animals were reared in large groups (gregarious state) at 28 °C with a 12 h / 12 h light / dark cycle; adult locusts from either sex were used for experiments. Limbs and wings were cut off, the animals were fixed on a metal holder with dental wax, and the head capsule was opened frontally, providing access to the brain. The esophagus was cut inside the head, close to the mandibles, and the abdomen’s end was cut off to take out the whole gut through this opening. The brain was freed of fat, trachea and muscle tissue and was stabilized with a small metal platform that was fixed to the head capsule. Shortly before recording, the brain sheath was removed at the target site with forceps, permitting penetration with sharp glass electrodes. The brain was kept moist with locust saline (Clements and May, 1974) throughout the experiment.

### 2.2 Intracellular recording and histology

Sharp microelectrodes were drawn with a Flaming/Brown filament puller (P-97; Sutter Instrument), their tips filled with Neurobiotin tracer (Vector Laboratories; 4 % in 1 mol· l^−1^ KCl) and their shanks filled with 1 mol· l^−1^ KCl. Intracellular recordings were amplified with a custom-built amplifier and digitized with a 1401plus (Cambridge Electronic Device, CED) analog-digital converter (ADC) or amplified with a BA-01X (npi electronic GmbH) and digitized with a Micro mkII with an ADC12 expansion unit (CED). Signals were monitored with a custom-built audio monitor and recorded with Spike2 (CED). Neurons were traced by electrically injecting Neurobiotin (∼ 1 nA positive current for several minutes). Each neuron presented in this study originates from a different specimen. Brains were dissected and immersed in fixative· (4 % paraformaldehyde, 0.25 % glutaraldehyde and 0.2 % saturated picric acid, diluted in 0.1 mol · l^−1^ phosphate buffered saline [PBS]) over night, followed by optional storage at 4°C in sodium phosphate buffer until further processing. Brains were rinsed in PBS (4 × 15 min) and incubated with Cy3-conjugated streptavidin (Dianova; 1:1,000 in PBS with 0.3 % Triton X-100 [PBT]) for 3 d at 4°C. After rinsing in PBT (2 × 30 min) and PBS (3 × 30 min), they were dehydrated in an ascending ethanol series (30 %, 50 %, 70 %, 90 %, 95 %, and 2 × 100 %, 15 min each) and cleared in a 1:1 solution of ethanol (100 %) and methyl salicylate for 20 min and in pure methyl salicylate for 35 min, to finally mount them in Permount (Fisher Scientific) between two coverslips. For anatomical analysis, brains were scanned with a confocal laser-scanning microscope (Leica TCS SP5; Leica Microsystems). Cy3 fluorescence was elicited with a diode pumped solid-state laser at 561 nm wavelength. The resulting image stacks were processed with Amira 6.5 (ThermoFisher Scientific, Waltham, MA) and Affinity Photo (Serif, Nottingham, UK). The chirality of some neurons could not be determined because multiple neurons of the same neuron class but on both brain sides were stained in these cases.

### 2.3 Experimental Design

We used two monitors (FT10TMB, 10”, 1024×768 px at 60 Hz, Faytech, Shenzhen, China) that were placed 12.7 cm apart on the left and right side of the animal. They were mounted vertically to present sinusoidal grayscale grating patterns (Figure 1A). The displays were covered with diffuser sheets to eliminate light polarization inherent to LCD monitors. The patterns were drawn on the inner center-square (15.35 cm edge length) of the displays, covering 62.3° of the visual field on each side. The monitor brightness amounted to 1.12 ·10^11^ photons cm^−2^ ·s^−1^ when displaying a black area and 7.09 · 10^13^ ·cm^−2^· s^−1^ when displaying a white area. Monitor brightness was measured using a digital spectrometer (USB2000; Ocean Optics) placed at the position of the locust head.

**Figure 1.**
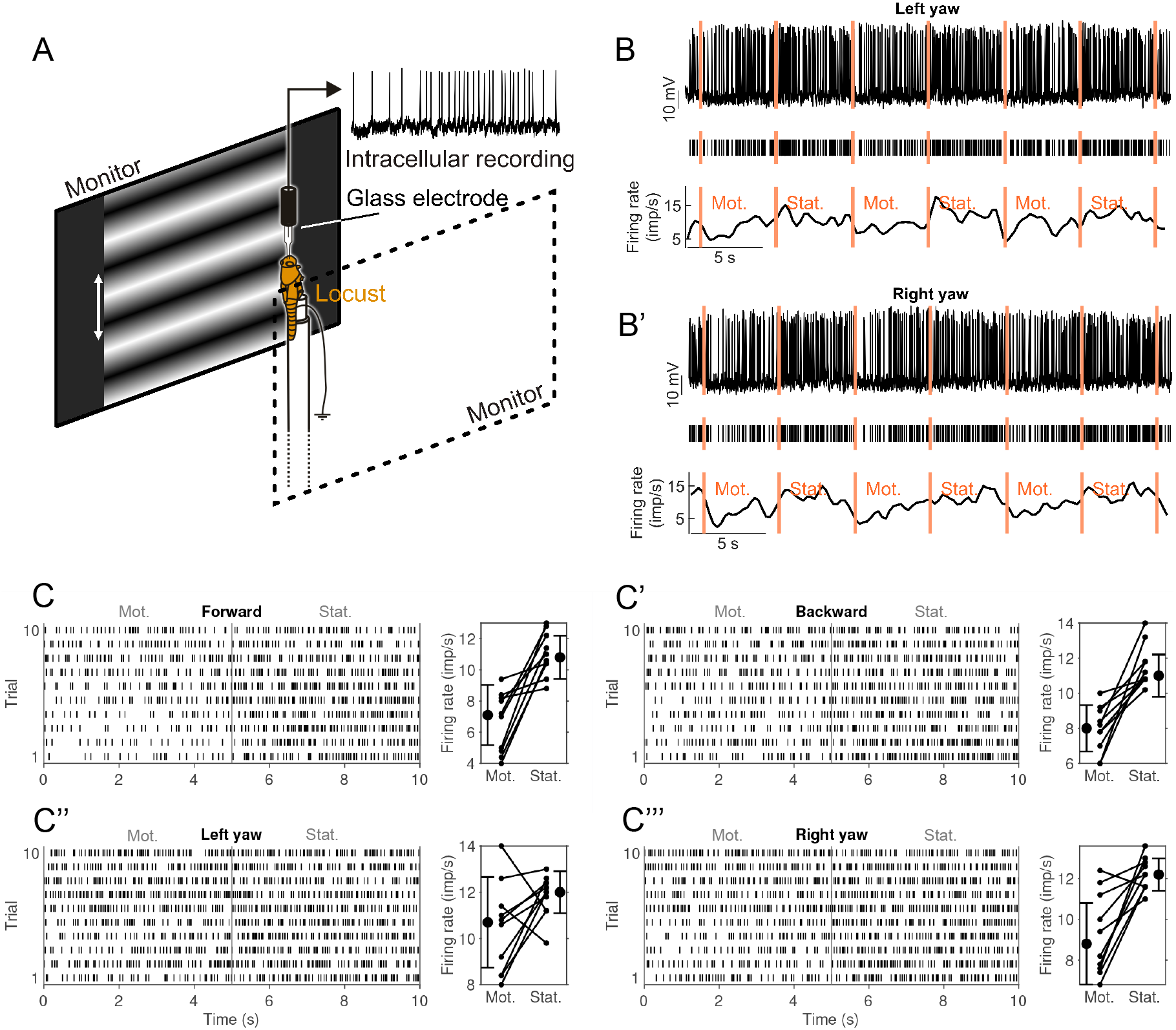
Experimental setup and visual-motion response of a CL1a neuron (neuron 550^*L*^ in Supplementary Figures 1 and 2). **(A)** Animals were mounted vertically and stimulated with motion of sinusoidal grating patterns on two laterally placed monitors. **(B)** Response of a CL1a neuron to wide-field visual motion that simulated horizontal left turning (left yaw). Raw data (top), detected spikes (middle) and smoothed firing rate estimate (bottom). Vertical lines indicate onset of stimulation phases: Motion (Mot.) and stationary phase (Stat.) were alternated, each pair constituting one stimulation trial. **(B’)** Same as B but for simulation of horizontal right turn motion (right yaw). **(C)** Raster plot (left) of all left-turn trials. Vertical line at 5 s indicates onset of stationary phase. Diagram on the right shows differences in firing rate between the motion (Mot) and stationary phase (Stat.) for each trial and mean firing rates for all trials. Error bars denote standard deviation. **(C’**,**C”**,**C”‘)** Same as C but for **(C’)** backward motion, **(C”)** left yaw and **(C”‘)** right yaw rotation.

The grating patterns were animated to simulate self-motion to the animal. We tested translational (forward and backward) motion, yaw rotation (left and right turning), lift (upward and downward), and roll (counter clockwise and clockwise). Throughout this study, these direction labels refer to simulated self-motion directions and not absolute motion of the displayed patterns. Thus, “forward motion” means that both monitors displayed a grating pattern with horizontal bands (perpendicular to the locust’s body axis, cf. Figure 1A) that continuously moved from top to bottom. For the sake of readability, we use “visual motion” for this wide-field visual motion stimulation, although this term includes diverse visual stimulation types, such as looming objects, small moving targets or full-panoramic optic flow, neither of which we presented to the animals.

Each motion direction was tested in a series of trials; each trial consisted of two phases, a motion phase and an immediately following stationary phase (Figure 1B,B‘). All phases in the same recording lasted for five or six seconds. Each series consisted of two to five trials; each trial was immediately followed by the next one, unless it was the last of the series. Neurons typically responded strongly to the pattern display switch between series. Therefore, each series of a given motion direction was preceded by an adaptation phase which was discarded; this phase was a single stationary phase of the same pattern used during the upcoming series, immediately followed by the first motion phase of the series. If the same motion direction was tested in more than one series, all trials were treated as if they belonged to the same series. Not all neurons could be tested for all motion directions due to recording instability.

A separate PC running MATLAB (R2019, MathWorks) with the Psychophysics toolbox (Brainard, 1997) was used to generate the grating patterns (Figure 1A). The sine gratings had a spatial resolution of 0.005 cycles· px^−1^ (one sine cycle spanned 200 px) and were shifted with 2 cycles· s^−1^ during the motion phases. This PC was USB-connected to an Arduino Uno (Arduino) via which TTL pulses were sent to the ADC, recorded at 500 Hz. These pulses indicated grating pattern animation and onset of stimulation phases. Two squares with 30 px edge length in the top left corner of each display were used to indicate the presented motion type by flashing them white: Each motion type was assigned a distinct number of flashes (20 ms duration) that were generated at the end of the adaptation phase of each series. Each square was covered by a photo diode that picked up the white flashes and whose signal was recorded by the ADC at 200 Hz. This allowed for encoding the motion type of each stimulation series in the data file. The generation of each rectangle flash was also recorded via the Arduino as a TTL rectangle pulse of the same duration, which allowed for measuring the precise timing of stimulus display by cross correlating diode signal and TTL signal.

### 2.4 Statistical Analysis

Spikes were detected by median filtering (500 ms window width) the voltage signal and applying a manually chosen threshold. Spikes and non-spikes (gaps) within 2 ms time bins were counted during the whole 5 s long interval of each trial of stimulation condition. We chose 2 ms time bins for this analysis because this is the approximate length of the refractory period of the neurons.

#### 2.4.1 Motion Sensitivity

We define motion sensitivity as a neuron’s property to have different firing rates during motion and stationary phases. We analyzed motion sensitivity for each tested neuron and motion direction by comparing the neuron’s firing rate during the motion phase with that during the previous stationary phase. Firing probabilities were computed by integrating prior knowledge about compass neuron activity in general and the condition-specific data from each neuron via Bayesian inference. For each neuron *n*, we computed a posterior over three different hypotheses: First, that the firing probability in 2 ms time bins during the motion phase *r*_*m*_ is lower than the firing probability *r*_*s*_ during the stationary phase, *H*(*r*_*m*_ < *r*_*s*_), second, that the firing probabilities are equal *H*(*r*_*m*_ == *r*_*s*_), or third, that *r*_*m*_ exceeds *r*_*s*_, *H*(*r*_*m*_ > *r*_*s*_). A high posterior for the first or third hypothesis would indicate motion sensitivity, while a high posterior for the second hypothesis would indicate that the neuron does not respond to the motion stimulation.

Using Bayes’ rule, we computed the posterior distribution *P* (*H*|*D*) over the three hypotheses *H* ∈{*H*(*r*_*m*_ < *r*_*s*_), *H*(*r*_*m*_ == *r*_*s*_), *H*(*r*_*m*_ > *r*_*s*_)} given the experimental data *D*, assuming an uniform hypothesis prior, a Bernoulli observation model and a joint Beta prior for the firing probabilities. This joint prior was restricted by the firing probability constraints expressed in each hypothesis, e.g. for *H*(*r*_*m*_ < *r*_*s*_), the probability *P* (*r*_*m*_ ≥ *r*_*s*_) = 0 etc. For details, see appendix 5.1.To summarize the information embedded in this posterior and to simplify comparison across multiple neurons, we computed a single motion sensitivity score (MSS) per neuron and motion direction (dir):

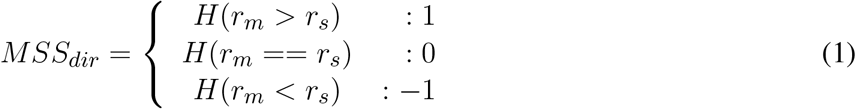

We weight this score with the corresponding hypothesis posterior probability and sum across all neurons of one type. The maximal value for one firing probability hypothesis is therefore equal to the number of neurons of a given type.

Further, we computed absolute motion sensitivity scores (AMSS) for four motion categories (cat), each comprised of two opposing motion directions *A* and *B*: translational motion (forward or backward direction), yaw rotation (left or right turning), lift (upward or downward), and roll (counterclockwise or clockwise):

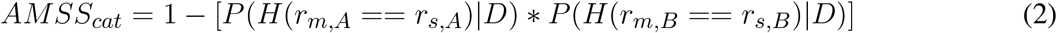

where *r*_*m,A*_ and *r*_*m,B*_ are firing probabilities during stimulation with opposing motion directions in the respective motion category. In other words, this score will be close to one if at least one motion direction of a category elicits a strong deviation from the stationary firing probability. We sum this score across all neurons of a given type.

#### 2.4.2 Direction Selectivity

We define direction selectivity as a neuron’s property to respond contrarily to two opposing motion directions *A* and *B*. We analyzed direction selectivity in the four motion categories outlined above: translation, yaw rotation, lift, and roll. In the following, the hypothesis *H*(*r*_*m,A*_ ≥ *r*_*s,A*_) = *H*(*r*_*m,A*_ > *r*_*s,A*_) ∨ *H*(*r*_*m,A*_ == *r*_*s,A*_) where ∨ indicates a logical or, and ∧ is a logical and.

We compute a direction sensitivity score as

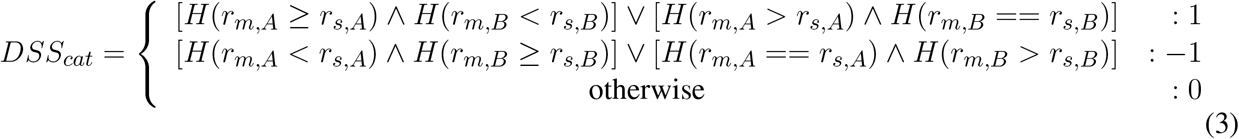

For example, *DSS*_*translation*_ is +1(−1) if the firing probability does not decrease during forward(backward) motion and decreases during backward(forward) motion, or if it increases during forward(backward) motion and does not change during backward (forward) motion. It is 0 if the firing probability changes in the same direction for both motion directions. We weight this score with the corresponding hypothesis posterior probability and sum across all neurons of one type. The maximal value for one firing probability hypothesis is therefore equal to the number of neurons of a given type, similar to *MSS*_*dir*_.

As an indicator for the total number of neurons with any direction sensitivity at all, we computed the expected absolute direction sensitivity score (ADSS):

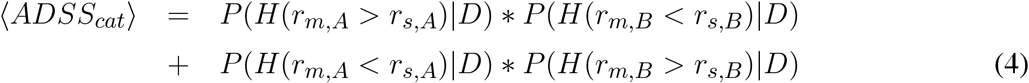

This score can take values between 0 and 1, with values close to zero indicating no direction selectivity and values close to one indicating direction selectivity, disregarding which motion direction elicits greater firing rates. We sum this score across all neurons of a given type.

The appendix 5.1 comprises a power analysis for the analyses of motion sensitivity and direction selectivity outlined above, indicating which difference in the recorded firing rates is considered evidence for the hypothesis that a neuron fires more in one of the two conditions.

### 2.5 Computational Model

All computations were performed with the Python programming language (version 3.10.8) and the Pandas (version 1.5.1) and PyTorch (version 1.13.0) libraries. Plots were created with the Matplotlib library (version 3.5.3).

Our model comprises CL1a and CL2 neurons, adopting the projection schemes proposed by Heinze and Homberg (2008). We assume that, as shown for E-PG and P-EN neurons in the fly (Turner-Evans et al., 2017), CL1a neurons provide synaptic inputs to CL2 neurons in the PB, which in turn provide synaptic inputs to CL1a neurons in the CBL. We further assume a combination of excitation and inhibition within the CL1a-CL2 connectivity instead of excitatory loops paired with global inhibition, as has been proposed for *Drosophila* (Turner-Evans et al., 2017). The firing rate neurons and synaptic connections in our model are linearized around their operating point, thus approximating their non-linear dynamics. We previously implemented this circuit with CL1a neurons exciting the CL2 neurons and CL2 neurons inhibiting the CL1a neurons (Pabst et al., 2022), referred to as Model A. In the present work we implemented an additional version of the model, referred to as Model B, where this relation is reversed because both versions are equally likely given the available data.

We represent the CL1a-CL2 connectivity with a matrix *M* - *M*_*A*_ and *M*_*B*_ for Model A and Model B, respectively. The matrix features additional self-recurrent connections at all neurons to enable the maintenance of a baseline activity. Weights are uniform for all excitatory and inhibitory connections, 0.5 and -0.5, respectively. The network’s activity is characterized by deviations from a baseline firing rate, represented by a vector *x*_*t*_ with components *x*_*t*,1:16_ and *x*_*t*,17:32_ covering the CL1a and CL2 neurons, respectively. The network is recurrent and iterated across time steps such that the activity at the next time step can be computed from the current activity:

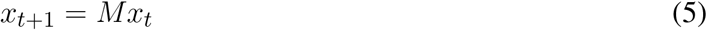

#### 2.5.1 Maintenance of a Stable Head Direction Encoding

In the framework outlined above, maintenance of the head direction representation or CL1a activity pattern *x*_1:16_ translates to an equality of *x*_*t*,1:16_ at time point *t* and *x*_*t*+1,1:16_ at the following time point, *t* + 1:

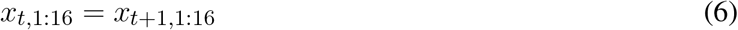

According to Equation 5, this is given if *Mx*_*t*_ = *x*_*t*_. We refer to such *x*_*t*_ as stable states. We defined CL1a activity targets 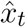 matching the tuning observed across the PB (Pegel et al., 2019; Zittrell et al., 2020) and employed an optimization algorithm to find stable states containing these targets. For more details, see Pabst et al. (2022).

#### 2.5.2 Rotation-induced Shifts of Compass Activity

We have previously described a possible computational mechanism that would produce a phasic shift from *x*_*t*_ to *x*_*t*+1_ with Model A (Pabst et al., 2022), representing the influence of rotational flow inputs on the compass system, putatively conveyed by TN or TB7 neurons. In the present work we adjusted the modulatory effect to allow for broad arborizations in the CBL, in addition to the PB, and optimized the synaptic weights to produce compass bump shifts with Models A and B. Furthermore, we applied additional constraints to both models for better alignment with physiological data: In Model A we found that purely feed-forward input to the CL1a and/or CL2 neurons cannot account for the observed shift behavior (Pabst et al., 2022). Instead, we found a modulatory mechanism that successfully shifts the compass bump. In our previous study we relaxed the original connectivity matrix *M*_*A*_ to allow for arborizations into adjacent PB columns and optimized the weights in this relaxed matrix 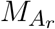 to achieve shifts to either direction, adding inhibitory or excitatory connections in up to two further PB columns on each side of all existing synapses. We constrained these shift-matrices 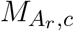 and 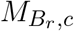 to better account for available physiological data: For the neurons modeled here, no arborizations broader than one column were found in the PB, while arborizations, in the CBL, especially in the upper layers, do in fact span three to five columns (Heinze and Homberg, 2008). We refer to these models as ‘relaxed and constrained models’.

## 3 RESULTS

We surveyed CX neurons at different integration stages for sensitivity to visually simulated self-motion (Figures 1,2). In total 62 morphologically identified neurons with arborizations in the CX were studied (Figure 2). These included 4 tangential input neurons (TL) to the CBL comprising the subtypes TL2 and TL3 (Figure 2A), 21 CL1a columnar neurons connecting the CBL to the PB, two CL2 columnar neurons connecting the PB, CBL and NO, five TB1 tangential neurons of the PB, three CPU1, seven CPU2 and one CPU5 neurons connecting distinct columns of the PB and CBU to the lateral complex (CPU1, CPU2) or a nodulus (CPU5), one CP1 and two CP2 neurons connecting the PB to distinct areas of the lateral complex (Figure 2B), eight PoU pontine neurons (Figure 2B), and various TU-type tangential neurons of the CBU (Figure 2A). We found sensitivity to visual self-motion in some neural classes while others did not respond to the stimulation.

**Figure 2.**
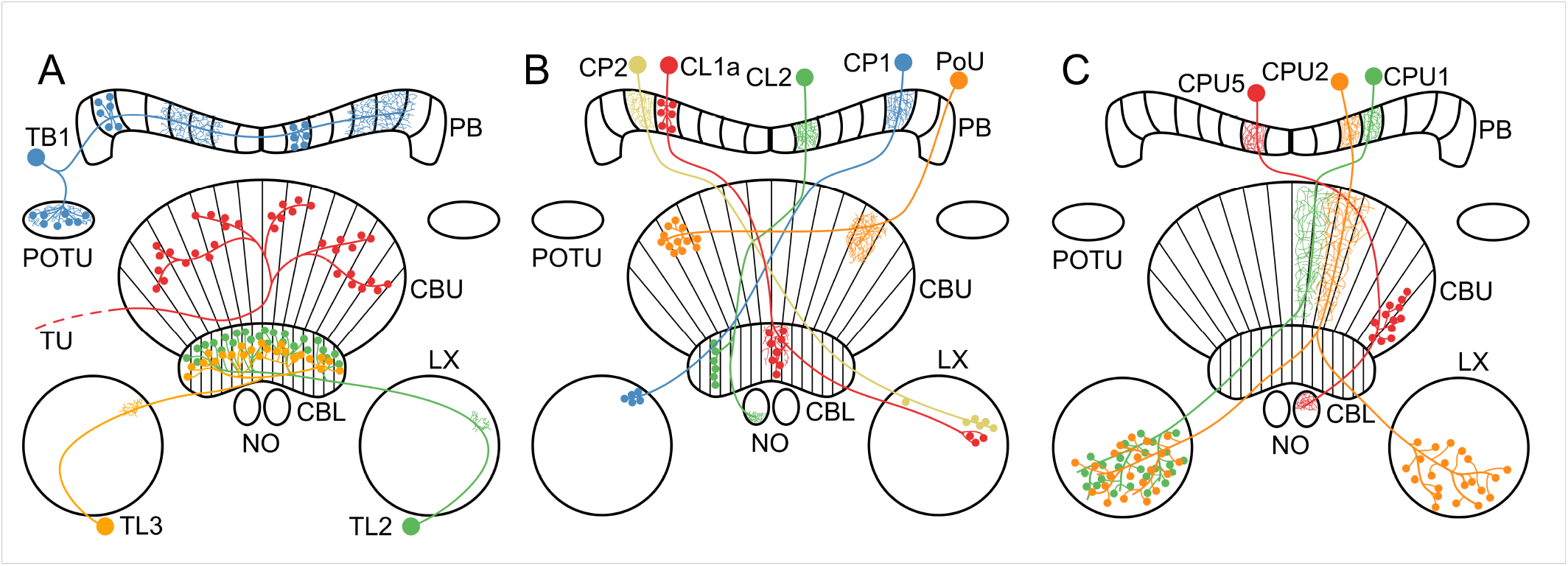
Morphology of neuron classes analyzed in this study. **(A–C)** Schematics of the locust central complex and associated neuropils (CBL, lower division of the central body; CBU, upper division of the central body; LX, lateral complex; NO, noduli; PB, protocerebral bridge; POTU, posterior optic tubercle) with individual neurons from different classes superimposed. Large dots indicate somata, small dots indicate axonal (presynaptic) arborizations, and fine lines indicate dendritic (postsynaptic) arborizations. Tangential neurons. We classified TU neurons as a group of diverse neurons that only have in common that they have large presynaptic arborizations in the CBU and input regions outside the central complex. Wiring schematics based on (von Hadeln et al., 2020). **(B**,**C)** Columnar neurons. Wiring schematics based on (Heinze and Homberg, 2008).

### 3.1 Visual Self-motion Sensitivity and Direction selectivity in the Central Complex

Neurons in most of the examined morphological classes shown in Figures 2A-C were not sensitive to the moving gratings. Some of the tested TL-, CL1a-, and CPU2 neurons, however, were sensitive to grating patterns moving in at least one motion direction (motion sensitivity; Figures 3A,3B). Response scores, indicating the sign of the firing rate change due to visual self-motion perception, were likewise inconsistently distributed within these neuron classes. Overall, within a given neuron class, individual neurons responded with excitation, inhibition or not at all to the same stimulus, independent of their brain side of origin (Figures 3A,3B). Two CL2 neurons, however, were not only motion sensitive but also responded differently to opposing motion directions (direction selectivity, Figures 3A,3B,4, and Supplementary Figure 2).

**Figure 3.**
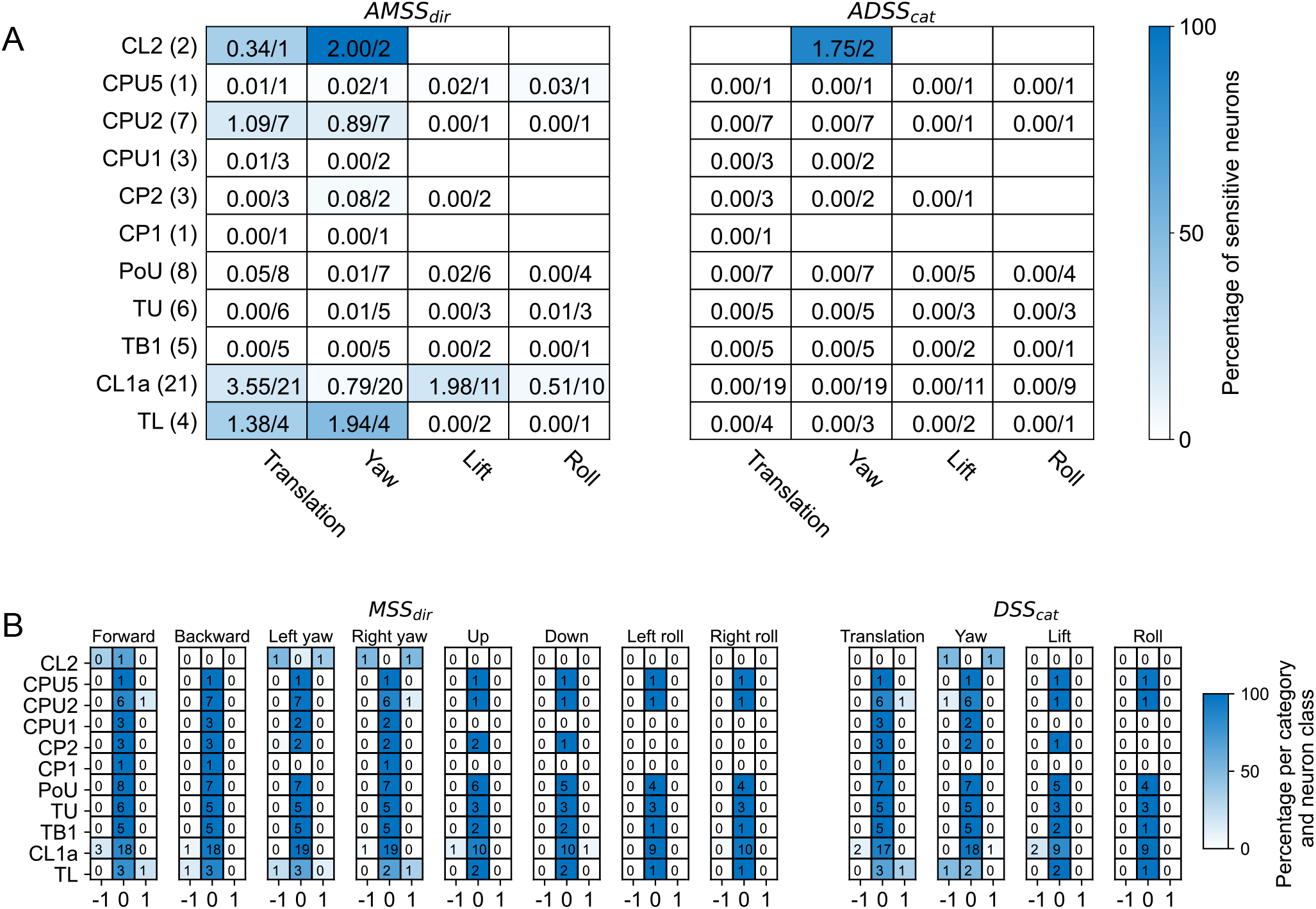
Overview of motion sensitivity and direction selectivity of all recorded neurons. **(A)** Absolute motion sensitivity scores per motion direction (*AMSS*_*dir*_, left) and absolute direction selectivity scores per motion direction category (*ADSS*_*cat*_, right), summed over neuron cell types. Absolute motion sensitivity scores take values between 0 and 1, with values close to 0 indicating no motion sensitivity and values close to 1 indicating motion selectivity, disregarding whether the neuron responds with an increase or decrease in activity. Absolute direction selectivity scores take values between 0 and 1, with values close to 0 indicating no direction selectivity and values close to 1 indicating direction selectivity, disregarding which motion direction elicits greater firing rates. Each cell holds the (rounded) sum of response scores over neuron cell types. Numbers are given as sums of scores over the total number of tested neurons. The fractions of summed scores and total possible scores are also indicated by the background color. The total number of recorded neurons for each neuron class is indicated in parentheses. Empty cells mean that no neuron was tested with the respective stimulus. **(B)** Distribution of motion sensitivity scores per motion direction (*MSS*_*dir*_, left) and direction selectivity scores per direction category (*DSS*_*cat*_, right), both per neuron class. Cell shading codes for the fraction of summed scores and total possible scores.

### 3.2 Yaw-rotation is processed by CL2 neurons

We recorded from two mirror-symmetric CL2 neurons. One neuron had smooth, presumably postsynaptic arborizations in the left NO and in column R4 of the right half of the PB, and beaded processes in layers 1-3 of column L2 in the left half of the CBL (Figure 4B). The second CL2 neuron had ramifications in the right NO, column L4 in left half of the PB, and column R2 in the right half of the CBL (Figure 4D). Both neurons were directionally selective for visual motion that simulated yaw rotation, but with opposite polarity (Figures 4A,A’,C,C’ and Supplementary Figure 2). The CL2 neuron with arborizations in the right half of the PB and in the left NO (unit 801^*R*^, Supplementary Figures 1 and 2) responded to right turns with an increase and to left turns with a decrease in firing rate, compared to baseline. The neuron was also weakly inhibited by forward motion. The CL2 neuron arborizing in the left half of the PB and the right NO (unit 800^*L*^ in Supplementary Figures 1 and 2) on on the other hand responded to left turns with an increase and to right turns with a decrease in firing rate. Responses to translational motion stimuli were not tested. CL2 neurons are part of the internal compass system in the locust CX (Pegel et al., 2018) and likely homologous neurons in *Drosophila* (P-EN) apparently signal rotational self-motion, updating the internal heading representation when the animal turns. Our data support the idea that the locust internal compass signal is also shifted during turns via asymmetric excitation and inhibition of CL2 neurons (Figure 5B‘). The site of this interaction may either be the NO (via TN neurons) or the PB (via TB7 neurons). Both cell types are, like their equivalents in *Drosophila*, the GLNO neurons and the SpsP neurons (Hulse et al., 2021) morphologically suited to provide asymmetric input to the CL2 population. Like in *Drosophila* P-EN neurons, the projections of locust CL2 neurons in the CBL are shifted by one column relative to the projections of CL1 neurons (Figures 5A,5B). A notable difference between compass representation in the locust and the *Drosophila* compass system is that the E-PG population activity peak in the EB results in two activity peaks with a fixed offset along the PB, while available data in the locust suggest a single peak along the PB that results from azimuthal tuning to celestial cues ((Pegel et al., 2019; Zittrell et al., 2020)). If so, locust CL2 neurons might have inhibitory connections to CL1a neurons (Figure 5B). However, these connections and their polarity are hypothetical as there are no data on functional connectivity in the locust CX. Alternatively, the observed tuning could be a consequence of the projection and connectivity patterns of CL1a and CL2 neurons.

**Figure 4.**
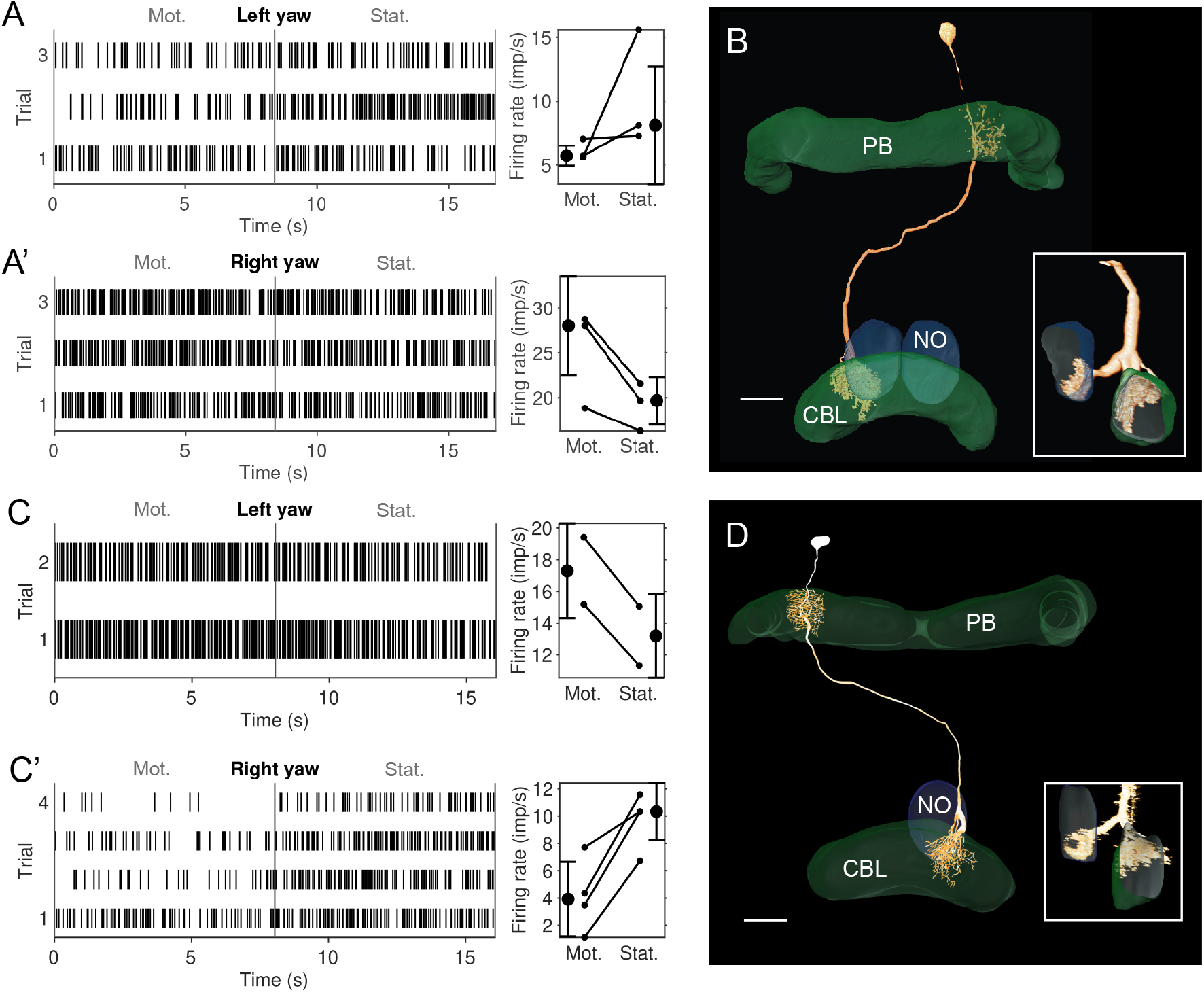
Physiological responses to yaw rotation and projections of CL2 neurons. **(A**,**A’)** Physiological response (raster plots and mean firing rates) to left yaw rotation (**A**) and right yaw rotation (**A’**) of the CL2 neuron shown in B (unit 801^*R*^ in Supplementary Figures 1 and 2). The neuron shows reduced firing rateduring simulated left yaw and increased firing activity during simulated right yaw. Vertical lines in the raster plots indicate onset of the stationary phase. **(B)** Skeleton view of the CL2 neuron (view from posterior) recorded in A and A’. The neurons arborized in column R4 of the right hemisphere of the protocerebral bridge (PB), layers 1-3 of column L2 in the CBL, and in the lower unit of the left NO. Inset shows sagittal view of ramifications in the lower division of the central body (CBL), and the left nodulus (NO). Scale bar: 40 µm. **(C**,**C’)** Raster plots and changes in firing rate during simulated yaw in the second CL2 neuron, shown in D (unit 800^*L*^ in Supplementary Figures 1 and 2). The neuron increased its firing rate during simulated left yaw (**C**) and decreased its firing rate during simulated right yaw (**C’**). **(D)** Two-dimensional reconstruction of the neuron from confocal image stacks (view from posterior). It arborized in column L4 of the left hemisphere of the PB, column R2 in the CBL, and in the lower unit of the right NO. Inset shows sagittal voltex view illustrating ramifications in the CBL and NO. Scale bar: 40 µm.

**Figure 5.**
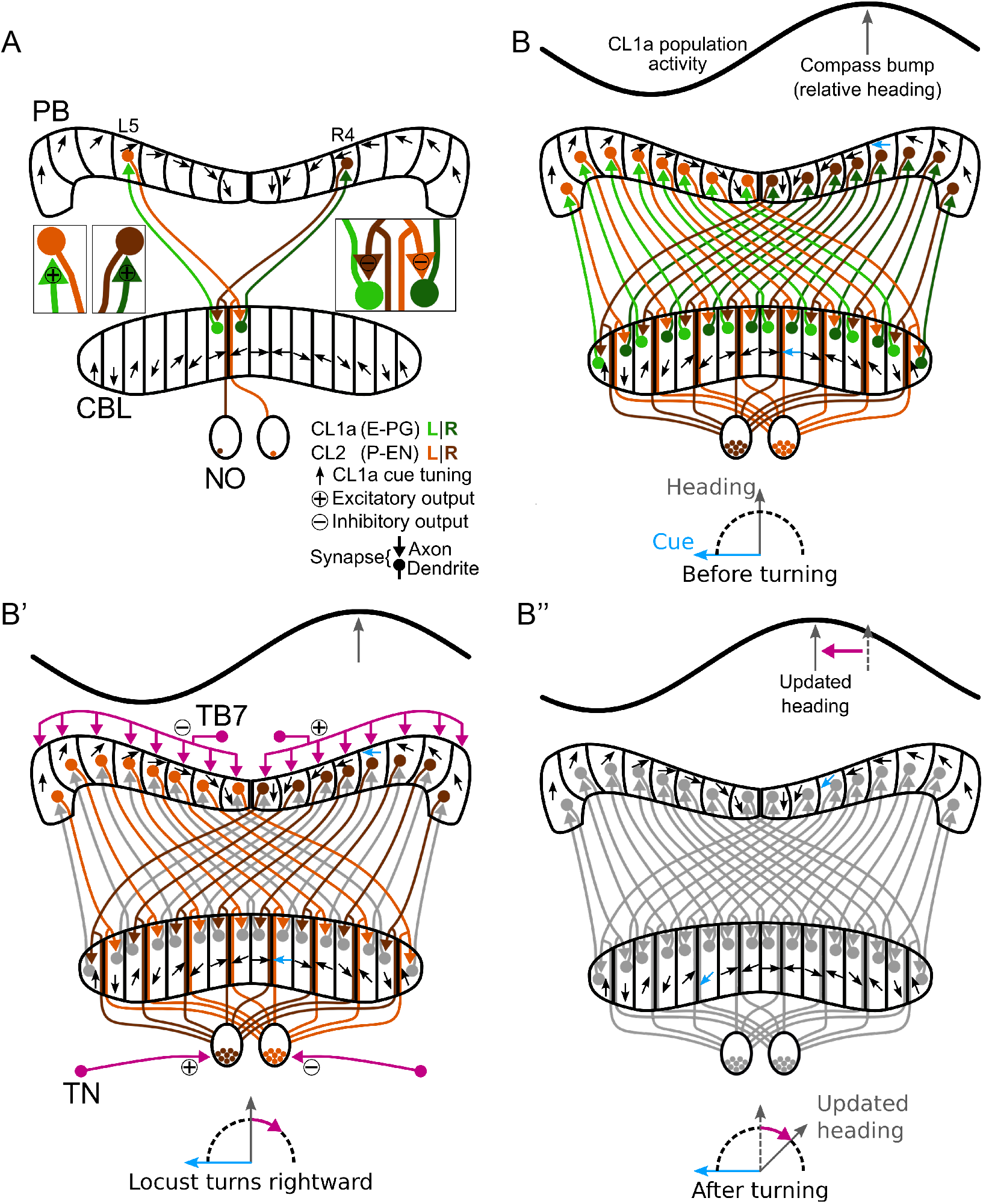
Schematic wiring diagram of CL1a and CL2 columnar neurons in the central complex and hypothetical shift mechanism of the internal heading signal in the PB. **(A)** Schematic wiring diagram of the CX with a subset of the involved neuron types: CL1a and CL2 neurons are connected to one another in the protocerebral bridge (PB) and lower division of the central body (CBL), while CL2 neurons also have postsynaptic arborizations in the noduli (NO). CL1a neurons are topographically tuned to solar azimuth along the PB (black arrows). **(B-B”)** Hypothetical shift mechanism of the internal heading signal in the PB. **(B)** Full population of CL1a and CL2 neurons and initial activity state in the network: With anenvironmental cue (sun) 90° left of the locust (bottom), the CL1a population activity (top) has a distinct maximum according to the neural tuning (highlighted arrows in PB and CBL). **(B’)** When the locust turns right, CL2 neurons are excited or inhibited depending on their brain side. Neurons that innervate the left (right) NO are excited (inhibited) by tangential neurons (TN) from the lateral complexes and relay onto CL1a neurons from the left (right) half of the PB. This asymmetric input may analogously be conveyed in the PB by tangential neurons (TB7) from the superior posterior slope. **(B”)** After turning, the CL1a population activity maximum is shifted so that the neural heading estimate accordingly represents the new heading relative to the external cue. Wiring schemes from (Heinze and Homberg, 2008), topographic tuning in the PB and CBL based on (Zittrell et al., 2020).

### 3.3 Computational Model

#### 3.3.1 Maintenance of a Stable Head Direction Encoding

Model A and B maintain an initial CL1a activity pattern with an activity maximum or compass bump representing head direction relative to a global cue, such as the sun, when no yaw rotation is simulated. The CL2 population’s activity is constrained by the polarity of synapses: We have previously shown that with CL1a neurons exciting CL2 neurons in the PB and CL2 neurons inhibiting CL1a neurons in the CBL in Model A, each CL2 neuron must have the same activity as the CL1a neuron associated with the same PB column to maintain a stable CL1a activity pattern (Pabst et al., 2022). In contrast, in Model B, where CL1a neurons inhibit CL2 neurons and CL2 neurons excite CL1a neurons instead, the CL2 activity pattern across the PB must be the inverse of the CL1a pattern for CL1a activity maintenance.

#### 3.3.2 Rotation-induced Shifts of Compass Activity

Feed-forward input to the CL1a/CL2 neurons can induce compass bump shifts neither in Model A nor in Model B. However, the weights in relaxed versions of the connectivity matrices 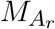 and 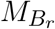 (Figures 6B,B‘) could be optimized to induce compass bump shifts in both versions of the model. Optimizations for both Model A and Model B resulted in a connectivity with excitatory synapses from CL1a onto CL2 neurons in the PB, a characteristic of Model A during compass bump maintenance (Figures 6B,B’). In the ‘relaxed and constrained models’, we allowed the optimizer to broaden arborizations of CL2 neurons in the CBL to more than one column and reduced arborizations of CL1a neurons in the PB to single columns (Heinze and Homberg, 2008). Compared to the results obtained with the ‘unconstrained relaxed models’, optimization converged at a solution where the weights for self-recurrent connections had greater absolute values (Figures 6C,C‘). Lastly, we enabled the addition of connections among CL2 neurons of the same hemisphere during the optimization process. Projection patterns suggest that all CL2 neurons of the same hemisphere branch in the lower unit of the contralateral NO, like P-EN neurons in *Drosophila* (Wolff et al., 2015) and the bumblebee (Sayre et al., 2021). Optimization converged at a solution with excitatory and inhibitory connections among CL2 neurons branching in the same nodulus. With these additional connections, synapses from CL2 onto CL1a neurons are weaker than in the other model versions (Figures 6D,D‘).

**Figure 6.**
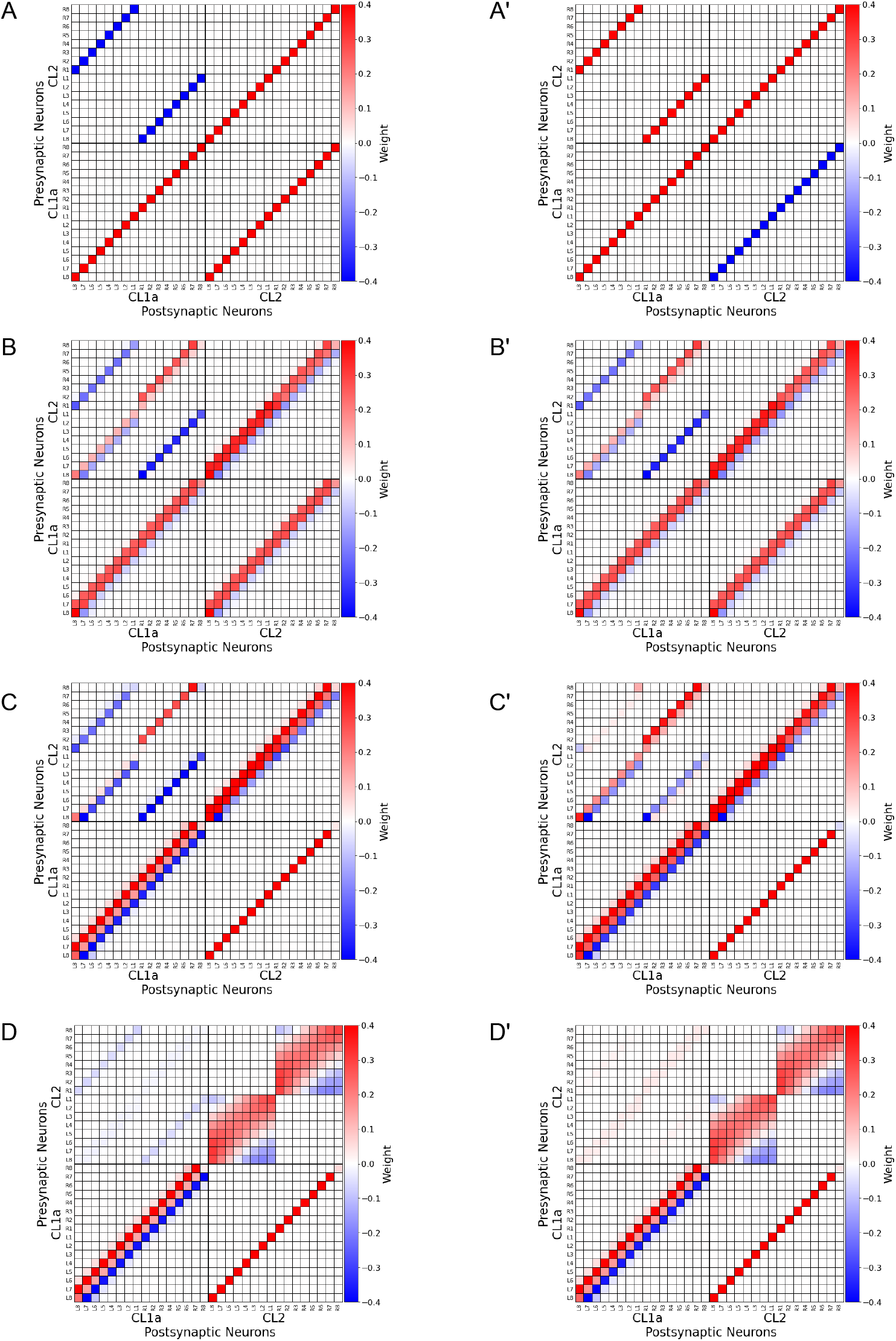
Computational Model. **(A-A’)** Connectivity matrices representing the projection and connectivity schemes shown in Figure 5B, with additional self-recurrent excitatory connections. Excitatory synapses are depicted in red, inhibitory synapses in blue. Neurons are indexed via the PB column (L8-R8) in which they arborize. **(A)** *M*_*A*_, implying excitatory synapses from CL1a onto CL2 neurons in the PB and inhibitory synapses from CL2 onto CL1a neurons in the CBL. **(A’)** *M*_*B*_ implying a reversed polarity. **(B-B’)** Relaxed versions 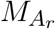 and 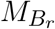 of the matrices shown in A-A’ optimized to produce shifts of the activity patterns **(C-C’)** Constrained versions 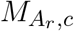 and 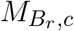 of 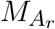 and 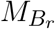, representing broader arborizations in the CBL but not the PB (compared to *M*_*A*_ and *M*_*B*_). **(D-D’)** 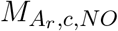 and 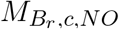; Same as C-C’ but with added connections among CL2 neurons branching in the same hemisphere of the PB and the same nodulus and optimized to shift activity patterns *x*.

## 4 DISCUSSION

We analyzed the sensitivity to visually simulated self-motion in different neuron classes in the locust CX network, from input-providing neurons (TL, TU neurons) to intermediate stage neurons (CL1a, CL2, POU, and TB1) and output neurons (CPU1, CPU2, CPU5, CP1, and CP2). Neurons in most of the investigated classes were not sensitive to visual self-motion. We hardly encountered consistent responses within the same neuron class, suggesting that single cells flexibly switch their cue sensitivity based on the internal state of the animal and environmental conditions. Exceptions were CL2 neurons, which mirror-symmetrically encoded yaw rotation direction, depending on the brain hemisphere in which they arborized, suggesting a role in keeping the internal compass system up to date during turning. A large fraction of cell types studied here (TL, CL1a, CL2, TB1, CPU1, CPU2, CP1, CP2) are elements of the sky compass system in the CX of the locust (Vitzthum et al., 2002; Heinze et al., 2009; Bockhorst and Homberg, 2015; Pegel et al., 2018; Zittrell et al., 2020). These neurons are sensitive to the azimuth of an unpolarized light spot (simulated sun) as well as to the polarization pattern above the animal (simulated sky) matching the position of the sun (Zittrell et al., 2020). The preference angles for solar azimuth in columnar neurons of the PB showed that solar azimuth is represented topographically across the columns of the PB as illustrated in Figure 5. The lack of responses to large-field motion stimuli in most of these neurons is in contrast to data from Rosner et al. (2019), who showed that a majority of sky compass neurons in the locust CX (types TL, CL1, TB1, CPU1, CPU2) were sensitive to progressive motion simulated through moving gratings. The reason for these different results most likely lies in different preparations of the animals. While in this study, legs and wings were removed, animals in the study of Rosner et al. (2019) had their legs attached and could perform walking motion on a slippery surface. Therefore, while the responses to sky compass signals may be affected only mildly, differences in behavioral context and internal state apparently play a major role for the sensitivity of sky compass neurons to visually simulated self-motion. Neurons of the CBU (PoU, TU, CPU5) that are not directly involved in sky compass signaling, were, likewise, unresponsive to visual self-motion. This coincides with studies on *Drosophila* that found that responsiveness of neurons of the fan-shaped body (corresponding to the locust CBU) to motion stimuli highly depended on whether the animals were actively engaged in flight (Weir and Dickinson, 2015; Shiozaki et al., 2020). It is therefore likely, as for neurons of the sky compass system, that neurons at this integration stage are silent in locusts under the constrained conditions of our experiments. H∆b neurons in *Drosophila* (corresponding to PoU neurons in the locust) integrate external and internal self-motion cues to transform egocentric directions into world-centric coordinates (Lu et al., 2022; Lyu et al., 2022). The lack of mechanosensory feedback under our experimental conditions likely explains why PoU neurons did not respond to purely visual self-motion cues. Under such conditions, PoU neurons, instead, strongly respond to looming objects (Rosner and Homberg, 2013), thus they might rather be involved in escape reactions when quiescence is signaled by the body. In general, physiological activity of locust CX neurons is considerably affected by active leg movement (Rosner et al., 2019). In our study, the legs were cut off, eliminating any proprioceptive sensory feedback.

In contrast to the lack of responsiveness in most cell types, two mirror-symmetric CL2 neurons showed robust responses to simulated yaw rotation with opposite directional preference. Inspired by the proposed role of P-EN neurons in *Drosophila* (corresponding to CL2 neurons in the locust) in updating and shifting the activity peak across the columns of the PB, we developed a computational model testing the likely function of CL2 neurons in the locust. The computational model of the CL1a-CL2 network resembles the recurrent loop connectivity between E-PG and P-EN neurons accounting for angular velocity integration in the *Drosophila* CX (Turner-Evans et al., 2017, 2020; Hulse et al., 2021). However, distinct differences exist, based on the 360° angular representation in the locust PB (Pegel et al., 2019; Zittrell et al., 2020) compared to the 2 × 360° representation of space in the *Drosophila* PB. While in *Drosophila* E-PG neurons form a 360° representation of space in the ellipsoid body, two opposite 180° representations of space would be topographically intercalated in the CBL of the locust (Figure 5A). In *Drosophila* P-EN and E-PG neurons are connected by recurrent excitatory loops with additional global inhibition (Turner-Evans et al., 2017). In the locust, instead, both inhibitory and excitatory connections between CL1a and CL2 neurons are required. Our model assumes homogeneously inhibitory or excitatory synapses from one neuron population onto the other. We implemented two versions of the same model, differing only in the polarity of CL1a-CL2 connections. Both versions proved suitable for compass state maintenance but required different activity patterns in the CL2 population. The version of the model with CL1a neurons inhibited by and exciting CL2 neurons requires the CL2 population activity to equal that of the CL1a population. The reversed version of this model in turn requires the CL2 population activity to be the inverse of the CL1a activity pattern. Physiological data revealing the relationship between the activities of these two populations would aid model evaluation and refinement. Close to equal E-PG and P-EN bump positions have been found in *Drosophila* moving at a low angular velocity, with an offset increasing with angular velocity (Turner-Evans et al., 2017). Neither of our versions could perform a shift of compass activity with a feed-forward input only, which might be due to the fact that our models do not include a closed loop from one end of the PB/CBL to the other. The inclusion of further neuron types might in fact close this gap and is the prospect of future work. CL1b-d neurons (Heinze and Homberg, 2008; Heinze et al., 2009) might, in addition, further stabilize the compass representation during standstill or forward motion. Franconville et al. (2018) reported that connections from E-PG onto P-EN neurons in the PB are mediated by ∆7 neurons. As TB1 and TB2 neurons cross the midline of the locust PB, they are, in addition to contralateral processes observed in some CL1 neurons innervating the innermost columns of the PB (Sayre et al., 2021), candidates for mediating ring closure. An internal compass representation must adapt to a new heading direction when the animal turns. In the CX, this is likely accomplished by integrating rotation cues of different modalities.

Two entry sites into the CX network for information on rotational self-motion have been proposed so far, based on work in the fruit fly: i) The PB, where neurons may receive asymmetric input excited depending on turning direction, conveyed via IbSpsP neurons (TB7 neurons in the locust) (Hulse et al., 2021). These neurons connect specifically to P-EN neurons (CL2 neurons in the locust). ii) The NO, where GLNO neurons (TN neurons in the locust) that receive input in the lateral complex and innervating one NO might be excited/inhibited depending on turning direction. P-EN neurons convey these asymmetric inputs to E-PG neurons via synapses in the ellipsoid body, leading to a shift of the internal heading representation according to turning (Green et al., 2017; Turner-Evans et al., 2017). We explored possible mechanisms inducing the compass bump shift on an algorithmic level.

Instead of an additive input, different modulations of the network connectivity can produce a shift of the compass network activity pattern. We adjusted the previously published modulatory effect (Pabst et al., 2022) to feature broader arborizations in both the PB and the CBL and repeated optimization with Models A and B. Relaxed versions (with broader arborizations in the PB and CBL) of both models could be optimized to shift the compass signal in both directions. Optimization of Model B converged at the same solution as optimization of Model A, with excitatory synapses from CL1a onto CL2 neurons in the PB, suggesting that Model A can better explain the behavior of the compass network. To obtain a better fit to physiological data, we narrowed down the width of arborizations in the CBL to three columns and we limited the arborizations in the PB to single columns. Again, optimization of Model A and B was successful and converged at modulated matrices with excitation in the PB in both cases. We further explored the possibility of synapses among CL2 neurons of the same hemisphere, which could occur in the lower units of the two NO and appear to be also present in *Drosophila* (Hulse et al., 2021). This additional degree of freedom rendered synapses from CL2 onto CL1a neurons mostly redundant for shifts. In contrast to the shift-inducing matrix presented previously (Pabst et al., 2022), the shift-inducing connectivity matrices shown here render a closed, ring-like architecture in the sky compass network. The linear model and discrete motion steps employed here are still quite abstract representations of the neuronal and behavioral characteristics of the locust. So far, our model is not dynamic; it switches between stable states but does not make the dynamics underlying the transitions explicit. We aim to increase the model’s biological plausibility by implementing velocity dependence in future work but expect the general principles of maintaining and updating the compass bump to hold independently of the level of analysis.

## AUTHOR CONTRIBUTIONS

FZ, RR, and UH designed the experiments, FZ, EC, UP and RR performed the experiments. FZ wrote manuscript. KP and DME designed the computational model and statistical analysis. KP revised the manuscript, analyzed the data and implemented the computational model with DME. DME and UH conceived, designed, and directed research, and helped write the manuscript. All authors contributed to the article and approved the submitted version.

## ACKNOWLEDGMENTS

We thank Stefanie Jahn for preparing Figure 4D and Martina Kern for maintaining locust cultures.

## DATA AVAILABILITY STATEMENT

The datasets analyzed and generated for this study along with the code written for analysis and modeling can be found in the data UMR repository (http://dx.doi.org/10.17192/fdr/76).

## 5 APPENDIX

### 5.1 Statistical Model and Power Analysis of Motion Sensitivity

We designed a Bayesian model for the evaluation of the experimental spiking data, to test the hypotheses that the firing probability of a motion phase *r*_*m*_ is smaller, equal or larger than the firing probability *r*_*s*_ during a stationary phase. We denote these hypotheses by *H*∈ {*H*(*r*_*m*_ < *r*_*s*_), *H*(*r*_*m*_ == *r*_*s*_), *H*(*r*_*m*_ > *r*_*s*_)}. Given the firing probabilities, we assume that the data *D* = (*s*_*m*_, *g*_*m*_, *s*_*s*_, *g*_*s*_) of one experiment, comprised of spikes *s*_*m*_, *s*_*s*_ during motion/stationary phases and corresponding non-spikes/gaps *g*_*m*_, *g*_*s*_, are generated by a Bernoulli process with a refractory period of 2 ms, which is typical for the neurons we investigate. There might be additional dependencies between spikes that are not captured by a refractory Bernoulli process, but these are not relevant for our hypotheses. The Bernoulli observation probability is given by

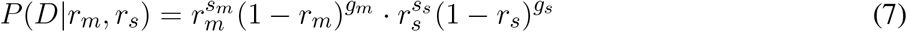

Since we are interested in hypotheses about firing probabilities relationships, we define a joint symmetric Beta prior with parameters *α, β* on *r*_*m*_ and *r*_*s*_, constrained by the hypothesis we wish to evaluate. We choose a symmetric prior to avoid a-priori biases beyond *H*. For *H*(*r*_*m*_ < *r*_*s*_), this prior is

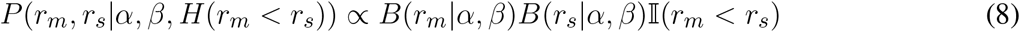

where *B*(*r*_*m*_ |*α, β*) is a Beta density in *r*_*m*_ and ǁ (*r*_*m*_ < *r*_*s*_) is an indicator function which is 1 if the condition in the parentheses is true, and 0 otherwise. This indicator function ensures that only hypothesis-conforming *r*_*m*_, *r*_*s*_ pairs have nonzero probability. The constant of proportionality can be obtained from the requirement that the prior be normalized. Thus, this prior can be written as

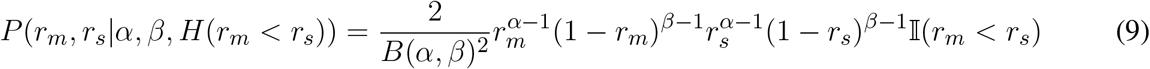

The prior resulting from *H*(*r*_*m*_ > *r*_*s*_) can be obtained by inversion of the < in the indicator function, whereas the prior for *H*(*r*_*m*_ == *r*_*s*_) is simply one Beta prior for both (equal) firing probabilities.

Since we are largely ignorant about the values of *α* and *β*, we chose these parameters by maximizing the differential entropy subject to the condition that the average firing probability is ≈0.05 in a 2 ms time bin, which is typical for our neurons. We found *α* = 0.96 and *β* = 18.28, and used these values for the rest of the analysis.

To compute the hypothesis posterior

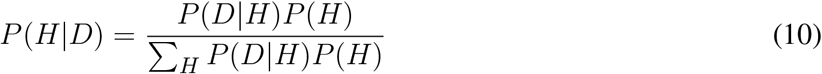

via Bayes’ rule, we chose a uniform hypothesis prior 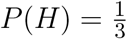. We evaluated the probability *P* (*D* |*H*) by marginalizing the firing probabilities using Equation 7 and Equation 9. For example, letting *H* = *H*(*r*_*m*_ < *r*_*s*_):

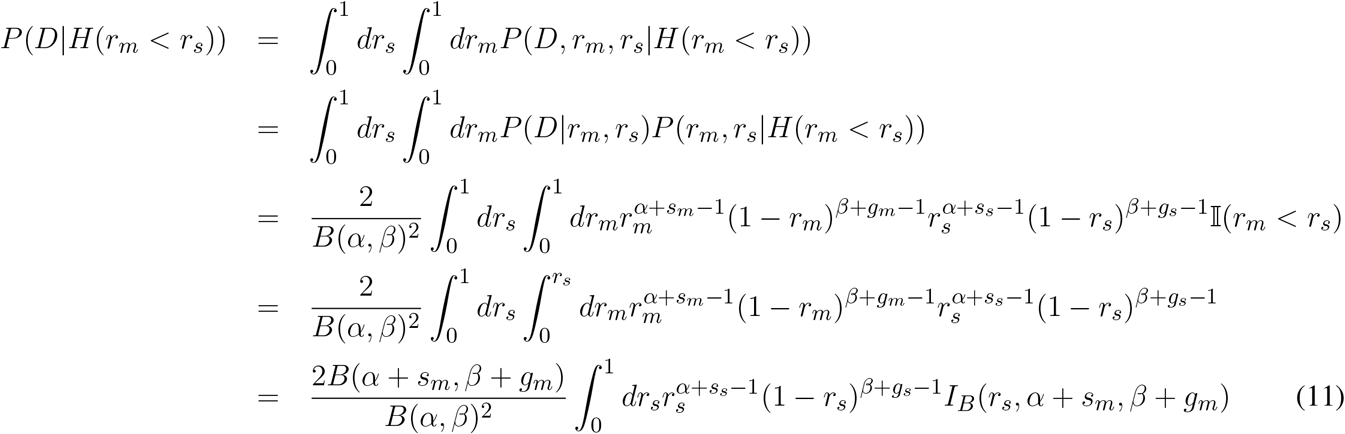

where *I*_*B*_(*r, α, β*) is an incomplete beta function in *r* with parameters *α, β*. We solved the last integral by Taylor-expanding *I*_*B*_(*r*_*s*_, *α* +*s*_*m*_, *β* +*g*_*m*_) to second order at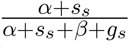, which yields a good approximation as long as *s*_*m*_ ≈ *s*_*s*_ and *g*_*m*_ ≈ *g*_*s*_. This is the case in our data.|

The probability *P* (*D* |*H*(*r*_*m*_ > *r*_*s*_)) can be evaluated by simply switching the roles of *r*_*m*_ and *r*_*s*_ in the above derivation. For *P* (*D*| *H*(*r*_*m*_ == *r*_*s*_)), where there is only one rate, the integrals can be solved analytically to yield the well-known result

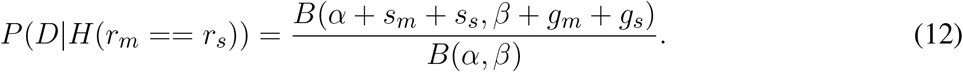

To facilitate interpretation of the values of the absolute motion sensitivity (AMSS, Equation 2) and the motion hypothesis posterior, which we use to average the motion sensitivity scores (MSS, Equation 1), we conducted a power analysis. We simulated 10,000 repetitions of a typical experiment in our study, where an animal is stimulated for 5 s with either stationary or motion input. We generated spikes according to the Bernoulli process assumption (Equation 7) with 2 ms time bins by drawing spike counts from a binomial distribution. The firing rate of the stationary phase was set to 25 Hz, which corresponds to a firing probability *r*_*s*_ = 0.05 and *N* = 2500 Bernoulli trials during a single run of the experiment. An experiment consisted of five simulated runs in the simulation. The firing probability during the motion phase was assumed to be a multiple of *r*_*s*_ in the range 1.15 … 1.30. This range is covered by a strongly responding neuron, see e.g. Figure 4A, right panel. To relate our motion sensitivity scores to standard measures used in statistical contexts, we evaluated the Bayes factor in favor of a changed firing rate during motion:

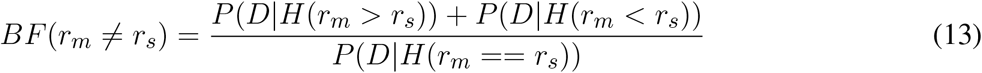

The simulation results are shown in Figure 7. The top panel shows the AMSS, the middle panel the corresponding Bayes factors. The dotted lines show the boundaries for weak and strong evidence according to Kass and Raftery (1995). For strong evidence, the firing rate ratio has to be greater than 1.25, which implies an average AMSS > 0.65. In the bottom panel, we plotted the hypothesis posterior, which we use for averaging of the MSS. Strong evidence for an increased firing rate (MSS=+1 in Figure 3A) requires MSS > 0.65.

**Figure 7.**
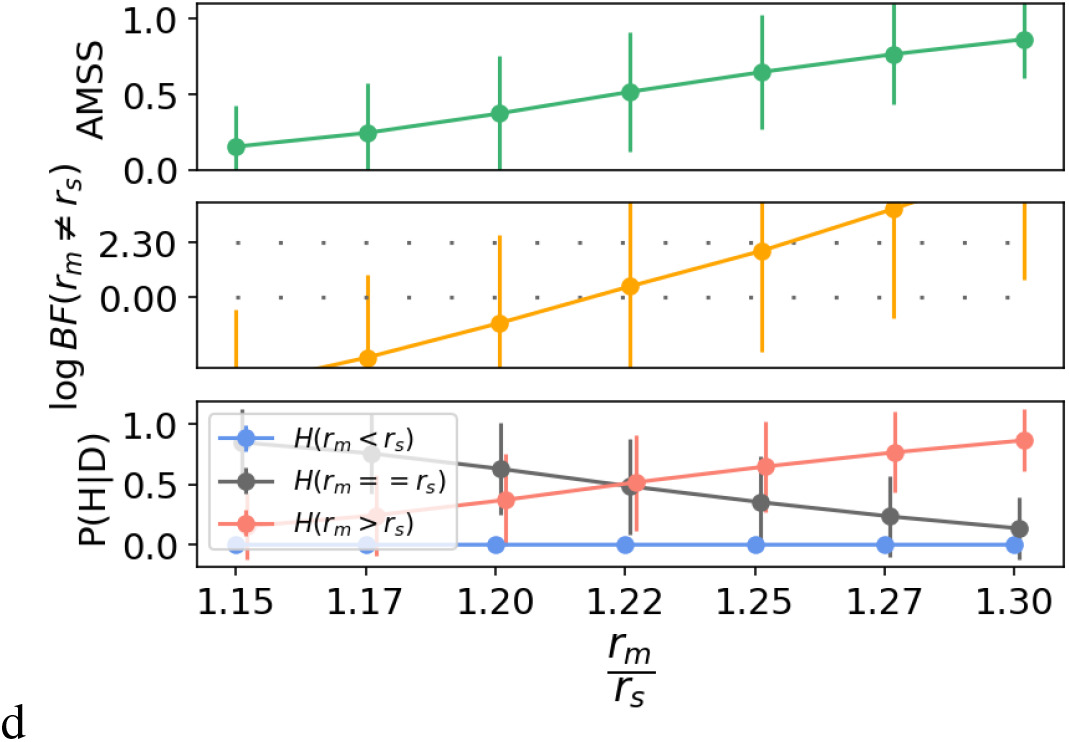
Power analysis of the Bayesian hypothesis comparison used for motion sensitivity analysis. The circles and error bars are means and standard deviations computed across 10,000 repetitions of a simulated experiment. The ratio of the motion phase firing rate *r*_*m*_ and *r*_*s*_ is shown along the abscissa. **Top**: absolute motion sensitivity (AMSS), cf. Equation 2. **Middle**: Bayes factor in favor of the hypothesis that the firing probabilities/rates are different during motion vs. equal rates, larger values represent stronger evidence. The dotted lines show the boundaries for weak and strong evidence according to Kass and Raftery (1995). **Bottom**: hypothesis posterior, used for the averaging of the motion sensitivity score (MSS), cf. Equation 1.The certainty of *H*(*r*_*m*_ > *r*_*s*_) increases with an increasing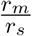ratio. 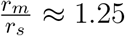 ≈ 1.25 is sufficient for strong evidence on average. For details, see text.

