## Supplementary material for "Integration of optic flow into the sky compass network in the brain of the desert locust"

### 1 SUPPLEMENTARY TABLES AND FIGURES

#### 1.1 Individual Motion Sensitivity Scores

We have described the motion sensitivity of a neuron by the posterior distribution over the hypotheses $H \in \{H(r_m < r_s), H(r_m == r_s), H(r_m > r_s)\}$ that the firing probability $r_m$ during motion stimulation is less/equal/more than the firing probability $r_s$ during stationary stimulation, see section 'Motion Sensitivity' of main paper. To summarize the information embedded in this posterior distribution per neuron, we computed the posterior expectation of the motion sensitivity score (MSS) per neuron and motion direction (dir):

$$\langle MSS_{dir} \rangle = P(H(r_m > r_s)|D) - P(H(r_m < r_s)|D) \tag{S1}$$

This score can take values between -1 and +1, with negative values indicating decreased firing and positive values indicating increased firing during the motion phase compared to the stationary phase. Furthermore, because this score represents a probability difference, $P(H(r_m > r_s)|D) \geq MSS_{dir}$. This implies that we can be at least as sure as $\langle MSS_{dir} \rangle$ of an increased motion response relative to the baseline, and conversely for negative scores. Values close to zero can result from equal activity during both phases (a high posterior for hypothesis $H(r_m == r_s)$) as well as from uncertainty about the three hypotheses (similar probabilities assigned to hypotheses $H(r_m < r_s)$ and $H(r_m > r_s)$ ). Therefore, we also computed an expected absolute motion sensitivity score (AMSS) for each direction:

$$\langle AMSS_{dir} \rangle = 1 - P(H(r_m == r_s)|D) \tag{S2}$$

This score can take values between 0 and +1, with values close to zero indicating no motion sensitivity and values close to one indicating motion sensitivity, irrespective of whether the motion response is a decrease or increase in firing compared to the stationary phase.

Supplementary Figure S1 shows the $\langle MSS_{dir} \rangle$ and the $\langle AMSS_{dir} \rangle$ for all neurons which we evaluated. The largest proportion of motion-sensitive neurons was found among CL1a, CL2, and TL neurons.

#### 1.2 Individual Direction Selectivity Scores

For a per-neuron summary of direction selectivity, we computed the posterior expectation of the direction sensitivity scores (DSS) for each motion category cat $\in$ {translation, yaw, lift, roll} (see section 'Direction Selectivity' of main paper):

$$\begin{aligned} \langle DSS_{cat} \rangle &= P(H(r_{m,A} > r_{s,A})|D) * P(H(r_{m,B} < r_{s,B})|D) \\ &- P(H(r_{m,A} < r_{s,A})|D) * P(H(r_{m,B} > r_{s,B})|D) \end{aligned} \tag{S3}$$

Here, $A, B$ are the opposite motion directions in each motion category:

| category | A | B |
|---|---|---|
| translation | forward | backward |
| yaw | left | right |
| lift | up | down |
| roll | left | right |

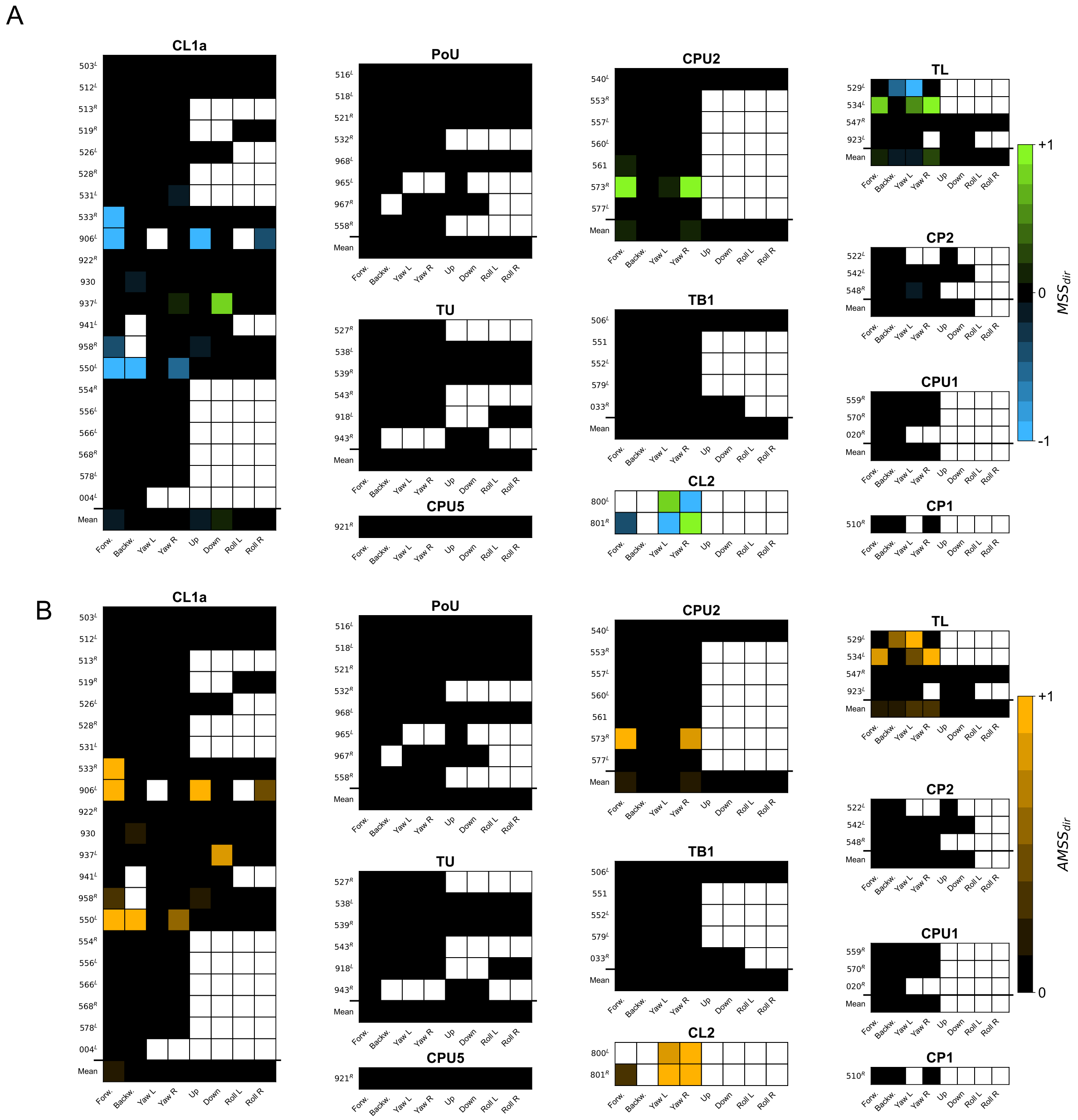


**Supplementary Figure S1.** Individual motion sensitivity scores per motion direction and neuron class. **(A)** False-color matrices show individual response scores for expected motion sensitivity $\langle MSS_{dir} \rangle$, indicating whether neurons responded to a given motion direction. Individual response scores range from -1 (strong inhibition compared to stationary stimulation, solid blue) to +1 (strong excitation, solid green) with 0 indicating a response that was best explained by $H(r_m == r_s)$, i.e. unchanged by motion (solid black). The 'mean' row, which was computed for more than four neurons of a type, holds the average response score per column, indicating whether neurons in the respective group tended to respond consistently. Rows are sorted for average response score; row numbers denote neuron ID, superscripts denote the brain side of the soma position. Empty fields indicate that the neuron was not tested with the corresponding stimulus; these fields did not contribute to the column average. **(B)** Same as A but for expected absolute motion sensitivity score $\langle AMSS_{dir} \rangle$, which ranges from +1 (strong firing rate change, solid orange) to 0 (no firing rate change, solid black). 'Mean' row has the same meaning as in **A**. Figure 1 of the main paper shows raw data of neuron $550^L$ (CL1a).

$\langle DSS_{cat} \rangle$ can take values between -1 and +1, with negative values indicating decreased firing and positive values indicating increased firing during the motion A phase compared to the motion B phase. Values close to zero can result from equal responses to both directions as well as from uncertainty about (meaning similar probabilities assigned to) hypotheses $H(r_m < r_s)$ and $H(r_m > r_s)$ indicating opposing responses. We therefore also computed the expected absolute direction sensitivity score (ADSS) per neuron, as an indicator for any firing rate changes between motions in opposing directions:

$$\begin{aligned} \langle ADSS_{cat} \rangle &= P(H(r_{m,A} > r_{s,A})|D) * P(H(r_{m,B} < r_{s,B})|D) \\ &+ P(H(r_{m,A} < r_{s,A})|D) * P(H(r_{m,B} > r_{s,B})|D) \end{aligned} \tag{S4}$$

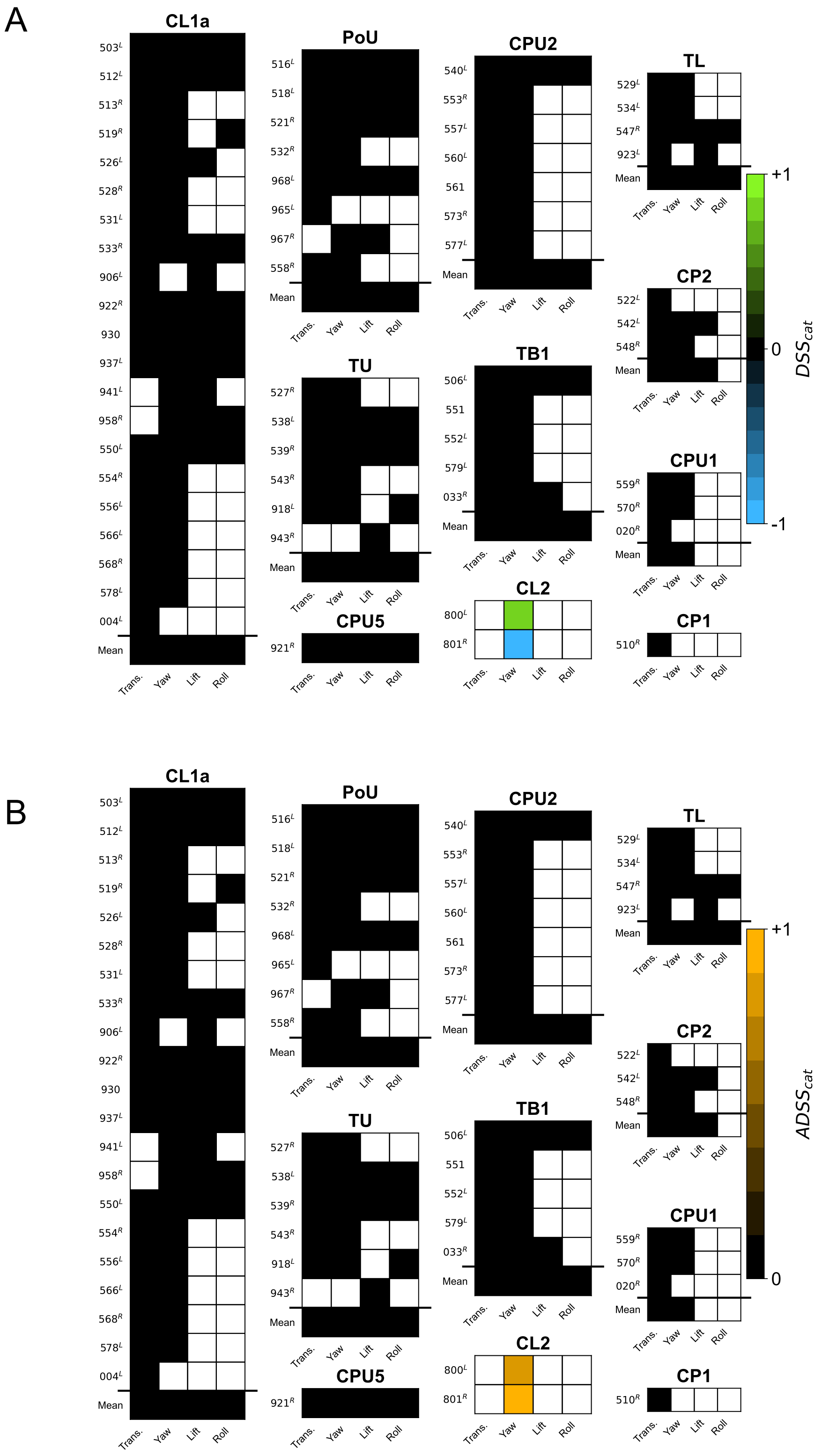


**Supplementary Figure S2.** Individual direction selectivity scores per direction category and neuron class. **(A,B)** Analogous to S1 but showing posterior expectations of direction selectivity scores $DSS_{cat}$ and $ADSS_{cat}$. The former indicates whether neurons responded by increased or decreased firing rate changes to opposing motion directions. The latter indicates whether the neuron responded to opposing motion directions at all.

Supplementary Figure S2 shows the $\langle DSS_{cat} \rangle$ and the $\langle ADSS_{cat} \rangle$ for all neurons which we evaluated. Only CL2 neurons exhibited direction selectivity.
